# Individual alpha frequency increases during a task but is unchanged by alpha-band flicker

**DOI:** 10.1101/559047

**Authors:** Michael J. Gray, Tatiana A. Emmanouil

## Abstract

Visual perception fluctuates in-synch with ongoing neural oscillations in the delta, theta, and alpha frequency bands of the human EEG. Supporting the relationship between alpha and perceptual sampling, recent work has demonstrated that variations in individual alpha frequency (IAF) correlate with the ability to discriminate one from two stimuli presented briefly in the same location. Other studies have found that after being presented with a flickering stimulus at alpha frequencies, perception of near-threshold stimuli fluctuates for a short time at the same frequency. Motivated by previous work, we were interested in whether this alpha entrainment involves shifts in IAF. While recording EEG, we tested whether two-flash discrimination (a behavioral correlate of IAF) can be influenced by ∼1s of rhythmic visual stimulation at two different alpha frequencies (8.3hz and 12.5hz). Speaking against the bottom-up malleability of IAF, we found no change in IAF during stimulation and no change in two-flash discrimination immediately afterwards. We also found synchronous activity that persisted after 12.5hz stimulation, which suggests that a separate source of alpha was entrained. Importantly, we replicated the correlation between IAF and two-flash discrimination in a no-stimulation condition, demonstrating the sensitivity of our behavioral measure. We additionally found that IAF increased during the task compared to rest, which demonstrates that IAF is influenced by top-down factors but is not involved in entrainment. In the framework of existing findings, we suggest that visual entrainment may involve ongoing perceptually-relevant oscillations from the delta to alpha frequency bands, serving to maintain rhythmic temporal expectations.

## Introduction

Numerous studies have shown that visual sensitivity fluctuates rhythmically. Depending on the task and methods used, rhythmic fluctuations in the detection of near-threshold stimuli have been observed in the delta (0.3hz-1hz; Fiebelkorn et al., 2011), theta (∼6-10hz; Busch et al., 2009; ∼4-10hz; Busch & VanRullen, 2010; 5-7hz; Benedotto et al., 2016), and alpha (∼10hz; Mathewson et al., 2009) frequency bands. More research in this domain has focused on alpha oscillations because of many earlier studies which have also demonstrated a strong relationship between alpha power and visual perception (e.g. Romei et al., 2008; Hanslmayr et al., 2007).

Several studies have suggested that the ongoing alpha oscillations which are involved in perceptual sampling can be entrained in a bottom-up manner to rhythmically flickering visual stimuli within the alpha frequency range (Mathewson et al., 2012; Spaak et al., 2014; Kizuk & Mathewson, 2016). In Spaak et al. (2014), a square stimulus was flickered at 10hz for 1.5s, causing the detection of a subsequently presented near-threshold stimulus to fluctuate at 10hz, in the same way that perception has been found to fluctuate spontaneously. Supporting the occurrence of the entrainment of ongoing alpha oscillations in the brain, they additionally found that 10hz neural activity persisted for about 300ms after cessation of the flicker.

It has been suggested that alpha entrainment involves a shift in an individual’s peak alpha frequency towards the stimulation frequency (IAF; Cecere et al., 2015; Hermann et al., 2016; Keitel et al., 2018). Cecere et al. (2015) found that the optimal timing of an illusion known as the sound-induced double-flash illusion was correlated with IAFs and that tACS stimulation at IAF ± 2hz caused changes in the optimal timing of the illusion that were consistent with entrainment-induced shifts in IAF. It remains an open question, however, whether rhythmic visual stimulation entrains alpha in the same way as non-invasive brain stimulation. Keitel et al. (2018) found no supporting evidence for a shift in IAF during quasi-rhythmic stimulation with average frequencies in the theta, alpha, and beta bands. In another recent study, it was found that very high intensity rhythmic stimulation at IAF and nearby frequencies produced stronger phase-locking compared to arrythmic stimulation but they did not examine whether shifts in IAF peaks occurred (Notbohm et al., 2016). So far, no studies employing rhythmic visual stimulation in the alpha-band have empirically tested whether alpha entrainment involves shifts in IAF and whether this corresponds to changes in perception.

With this in mind, our study sought to find behavioral and electrophysiological evidence for flicker-induced shifts in IAF by employing a task which was recently shown to correlate with IAF (Samaha & Postle, 2015). Samaha and Postle (2015) found that participants with faster IAFs were better able to discriminate one from two circles flashed briefly in the same parafoveal location. This suggested that those individuals were sampling the stimuli at a faster rate and were better able to segregate two-flash stimuli. We used this paradigm to test whether ∼1s of rhythmic stimulation at 8.3hz or 12.5hz would induce changes in participants’ ability to discriminate one from two flashes while concurrently measuring EEG. We predicted that 8.3hz stimulation would impair performance and decrease IAFs while 12.5hz stimulation would improve performance and increase IAFs. We also examined the EEG data for stimulation-induced synchrony after the offset of the flickering stimulus, which is frequently used as evidence for alpha entrainment (Mathewson et al., 2012; Spaak et al., 2014; Kizuk & Mathewson, 2016). Lastly, because it has also been shown that shifts in IAF and single-trial alpha frequency can occur due to top-down factors such as task demands (Haegens et al., 2014; Wutz et al., 2018), we also measured whether IAF changed from rest to the task.

Contrary to our hypothesis, we found that two-flash discrimination performance was not influenced by alpha-band flicker and IAFs during stimulation were unchanged. We additionally observed post-stimulation synchrony after 12.5hz flicker, suggesting that alpha entrainment occurred but did not involve IAF. Importantly, we also replicated the finding that IAF was correlated with individuals’ two-flash fusion thresholds (Samaha & Postle, 2015), demonstrating the sensitivity of our measure to reflect variations in IAF. We further found that IAF was increased during the task compared to an eyes-closed resting session, illustrating that although shifts in IAF are resistant to bottom-up influences, they do occur as a result of top-down factors.

## Methods

### Participants

Thirty-six undergraduate students participated in this study and were compensated with course credit (mean age: 20.34, SD: 3.13; 15 females, 5 left-handed). One participant did not complete our post-experiment questionnaire and thus was not included in these demographics. Including set-up, breaks, and clean-up, participants volunteered up to 4 hours of their time. The task itself lasted approximately 75 minutes. All participants self-reported having normal or corrected-to-normal vision and no history of neurological illness.

### Experimental Design

We tested whether a short period of visual flicker (8.3hz or 12.5hz) would influence participants’ ability to subsequently discriminate between one and two briefly flashed circles (i.e. the two-flash fusion task, previously shown to correlate with individual peak alpha frequencies (IAFs), Samaha & Postle, 2015). There was also a no-stimulation condition to replicate Samaha & Postle, 2015 and a stimulation condition in which the entrainer was stationary to test whether visual stimulation without any rhythmicity would influence two-flash discrimination. See Figure 1 for a schematic of the task.

**Figure 1.**
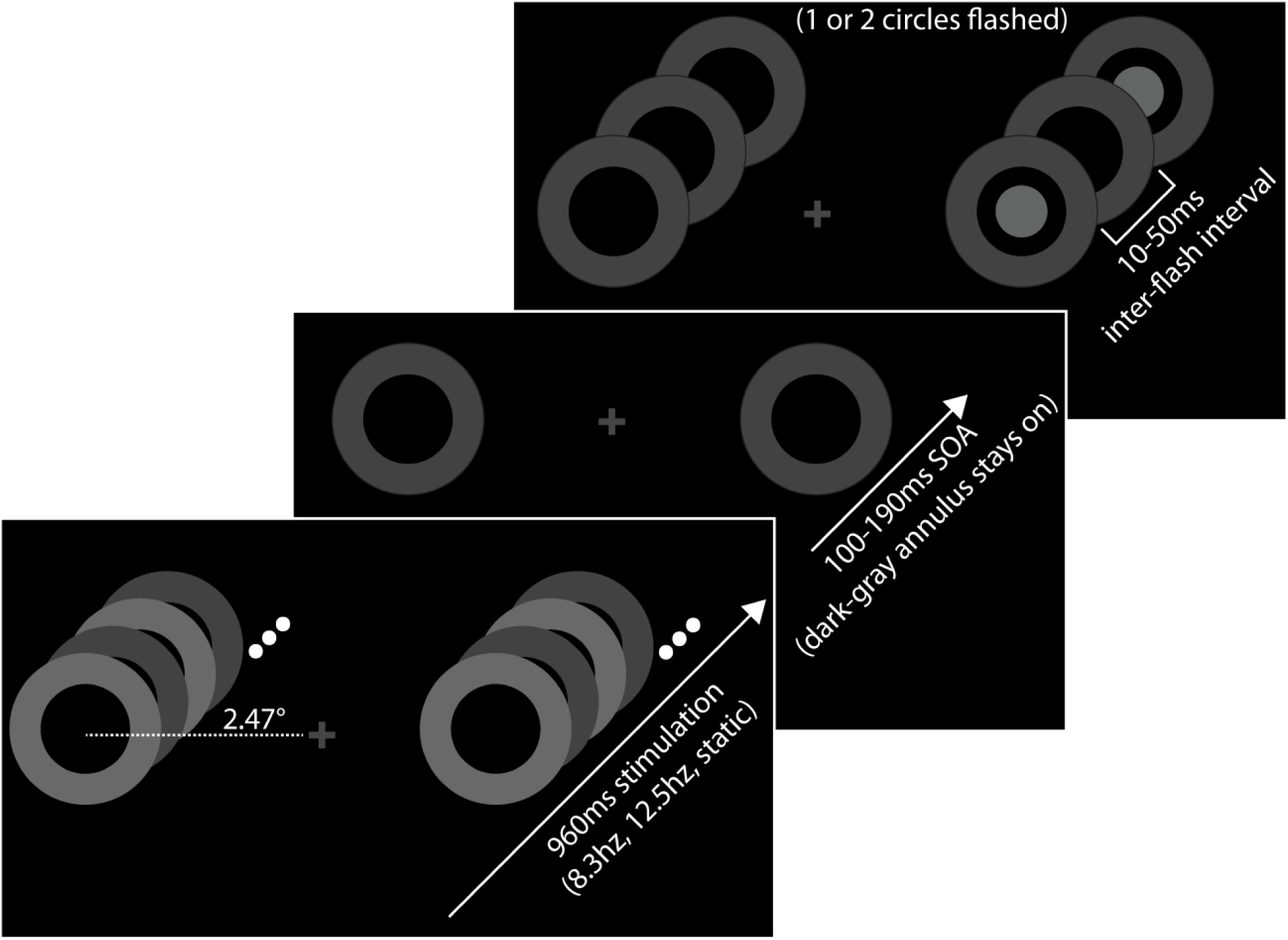
Task schematic. Stimuli are to-scale with respect to each other. Participants fixated on the central cross and after an inter-trial interval period (1800-2300ms), the annuli appeared and flickered between light-gray and dark-gray as shown. Following a brief SOA, one or two circles were flashed in the same location, within one of the annuli. Participants then responded via mouse click whether they perceived one or two flashes.

Participants were seated in a dimly lit experiment booth, 80cm from a 100hz CRT monitor (brand: iiyama; model: MA203DTA) set to 800 × 600 resolution. Before the main task, two 4-minute EEG recordings of resting alpha activity were obtained, one with eyes-open and another with eyes-closed. In the eyes-open recordings, participants were instructed to maintain their gaze on a fixation cross at the center of the screen. In the eyes-closed recordings, participants were told to sit still with their eyes closed. The order of these 2 sessions was counterbalanced across subjects.

Each trial in the main task proceeded as follows: Participants first fixated a light gray central cross (RGB = 70,70,70; 0.66° × 0.66°) on a black background for 1800-2300ms. Then, except in the no-stimulation condition, a light-gray annulus (RGB = 105,105,105; 3.74° × 3.74°) appeared 2.47° to the left and right of the fixation cross. In the 8.3hz and 12.5hz flicker conditions, the annuli alternated between light-gray and dark-gray (RGB = 65,65,65) every 60ms or 40ms, respectively, for 960ms. Note that this is of similar duration to existing studies which have found fluctuations in perception due to alpha-band flicker (560ms: Mathewson et al., 2012; 1500ms: Spaak et al., 2014; 560ms: Kizuk & Mathewson, 2016). In the 8.3hz condition, the annulus flickered for 8 cycles (120ms per cycle), while in the 12.5hz condition, the annulus flickered for 12 cycles (80ms per cycle). At the end of the last flicker cycle, the dark gray annuli remained on the screen to prevent masking effects as well as to reduce stimulus offset artifacts in the EEG. The final light gray entrainers changed to dark-gray at 900ms in the 8.3hz flicker condition and 920ms in the 12.5hz flicker condition in order to have a completed stimulation cycle after 960ms in both conditions. In the stationary stimulation condition the annuli remained light-gray for 910ms and then turned dark-gray. In the no-stimulation condition the single/double-flash stimuli were presented after the inter-trial interval (ITI) period.

After the last stimulation cycle in the stimulation conditions, the single/double-flash stimuli were presented, 100-190ms (10 sample points, one every 10ms) after the offset of the final light-gray entrainer. This range of stimulus offset asynchronies (SOA) ensured that there was an in-phase and an out-of-phase time point for both entrainment rhythms. Note that the abbreviation SOA is being used unconventionally in this study. SOAs were randomized and counterbalanced such that each stimulation condition had 20 trials at each SOA.

The timing and proportions of the single/double-flash stimuli were the same as Samaha & Postle, 2015. The circle used for the flash stimuli was 1.23° × 1.23°, RGB = 100,100,100. There were an equal number of single and double-flash trials and the flash stimulus occurred with equal probability, randomly on one side of the screen, within one of the dark gray annuli (which remained on the screen after the stimulation period). In double-flash trials, each flash was 40ms in duration, with an inter-flash-interval of 10-50ms (10ms increments). The durations of single-flash stimuli were equated to that of the two-flash stimulus events, such that single-flashes lasted between 90-130ms. Because the two-flash stimuli appeared within the previously-flickering annuli, a potential concern is that metacontrast masking could occur. This is not likely to be a confound because the SOAs used were longer than what would produce masking (Boyer & Ro, 2007).

Participants were told that there would be an equal number of one and two-flashes, and that the duration of the stimulus was not indicative of whether one or two-flashes had occurred. After the one/two-flash stimulus occurred, there was an 800ms delay before participants were cued by a slight brightening of the fixation cross (RGB = 90,90,90) to make their discrimination response. Participants made their responses with a mouse (left button = single-flash, right button = double-flash). Once a response was detected, the fixation cross darkened again to indicate the start of the next ITI.

There were 240 trials per condition, 40 of which were trials without single/double-flash stimuli after the stimulation period. Thus, there were 100 single-flash and 100 double-flash trials per stimulation condition, and 20 double-flash trials per inter-flash interval (10-50ms). The experiment was divided into 12 blocks, 3 blocks per stimulation condition. A break screen followed each block and participants could continue with the next block at their own pace. The order of the stimulation blocks was random with the constraint that no stimulation-type could repeat. Additionally, to help reduce differences in fatigue effects between conditions, the last 4 blocks always contained all 4 conditions in random order.

### Psychometric Function Fitting

The logistic function was used to compute psychometric curves and two-flash fusion thresholds for each participant using the PAL_PFML_Fit function in the Palamedes toolbox (Prinz & Kingdom, 2009). This function uses the Nelder-mead search algorithm to iteratively find the best-fitting curve according to 4 parameters: threshold (α), slope (β), guess-rate (γ), and lapse-rate (λ). Threshold, slope, and lapse-rate were free parameters and guess-rate was fixed at 0.5. Following the parameters used by Samaha and Postle, (2015), lapse-rate was bounded between 0 and 0.06. Goodness of fit measures were then computed using the PAL_PFML_GoodnessOfFit function. This function was used to calculate deviances for the actual fit and for 1000 fits simulated using a saturated model. This was done for each participant and each stimulation condition. The average proportion of deviance for all fits, pooled across stimulation conditions, was 0.70; SD = 0.27. Participants were excluded from a given condition when the deviance of the fit for that condition was unacceptable (exceeded the 95^th^ percentile of the deviances of their simulated fits) or the threshold for the condition was exceedingly high (more than 240 ms or more than 2.5 standard deviations of the mean threshold from the condition). Overall, we excluded 4 participants from the no stimulation condition, 2 participants from the stationary condition, 3 participants from the 8.3hz condition and 3 participants from the 12.5hz condition. Thus, only 11 (7.6%) of 144 fusion thresholds were excluded from further analysis.

### Post-Experiment Questionnaire

As an exploratory measure, we had participants complete a short questionnaire after the experiment. We were interested in whether IAF had any relationship to different short and longer-term traits relating to fatigue and concentration. On a scale of 1-10, we had participants rate perceived task difficulty, fatigue at the beginning of the task, fatigue at the end of the task, and whether they ever have difficulty concentrating. We also asked how many hours they slept the previous night and how many hours they usually sleep. Finally, the questionnaire also included 3 items about hobbies; in particular we asked about frequency of video game playing and expertise with dance and music.

### EEG Acquisition and Preprocessing

EEG data was sampled at 1000hz using a Neuroscan 64-channel EEG cap. Channels are arranged in this cap according to the extended 10/20 system. All data were filtered online with high-pass and low-pass filters set at 0.1hz and 100hz, respectively. The data were referenced online using electrode Cz, and then re-referenced offline to an averaged reference. Offline EEG data processing was done in MATLAB using custom scripts incorporating functions from the FieldTrip analysis toolbox (Oostenveld et al., 2011). Before epoching the data, bad channels were identified using statistics of neighboring channels and interpolated using spherical-spline interpolation. Re-referencing occurred after this step.

The eyes-closed and eyes-opened resting alpha recordings were each epoched into 240 1-s segments. The stimulation conditions in the main task were 960ms longer than the no-stimulation condition. Thus, for the stimulation conditions, data were epoched into 4s segments which included 1.5s pre-stimulation onset and 2.5s post-stimulation onset. Trials in the no-stimulation condition were epoched into 2.75s segments, 1.5s before and 1.25s after the onset of the flash discrimination stimulus.

A high-pass filter at 0.75 Hz was applied to each epoch using a fifth-order Butterworth filter with zero phase shift. Epochs were also linearly detrended and the main task data was additionally demeaned to the last 200ms before visual stimulation. Muscle artifacts were identified by band-pass filtering the data from 60 to 90 Hz with a fifth-order Butterworth filter. This data was then z-transformed and trials exceeding a z-value of 20 were removed. The data were then low-pass filtered at 40 Hz using a sixth-order Butterworth filter with zero phase shift and trials with amplitude fluctuations exceeding 75 uV over the posterior 1/3 of the cap were also removed. These procedures resulted in 10.1% of trials being rejected in the main task and 4.06% of trials being rejected in the resting alpha sessions. Lastly, the independent component analysis (ICA) function in FieldTrip was used to identify and remove ocular and heartbeat artifacts. The data were decomposed into 64 orthogonal components and were visually represented by topographies and time series. The components were inspected manually and those which were definitively an ocular or heartbeat artifact were removed.

### Individual alpha frequency (IAF)

IAF was computed for the eyes-closed and eyes-open resting alpha sessions, as well as for the task ITI, by finding the frequency with the local maximum in alpha power (8-13hz) for each subject at the sensor with greatest alpha power on the group level (PO6 for both resting sessions and for the task ITI). Each 1s epoch was first windowed with a Hamming window to prevent edge artifacts, zero padded to 10s to increase frequency resolution, and then submitted to a fast-Fourier transform (FFT). Power spectra were then computed by averaging across trials and then the local maximum between 8-13hz was found. The final 1s of the ITI was used to compare with resting IAFs. Spearman correlations were used for all IAF correlations to reduce sensitivity to outliers (as was done in Samaha & Postle, 2015). One participant did not show a peak in the ITI of the no-stimulation condition, so their data was excluded in this condition only.

To examine whether stimulation influenced IAF, we also computed IAFs for the last 500ms of the stimulation period using the same specifications as described above. In this analysis, more subjects did not show clear IAF peaks because power was much lower due to stimulus-evoked alpha desynchronization (e.g. Pfurtscheller & Aranibar, 1977). In addition to excluding data which showed no defined peak in the alpha range, we also manually excluded 2 participants in the 8.3hz condition and 1 participant in the 12.5hz condition who showed small stimulus-evoked peaks which were larger than their IAF peaks. In total, 10 of 36 subjects had at least one condition which had to be excluded. The 26 remaining subjects were included in a repeated measures ANOVA analyzing the effect of stimulation on IAF. In reporting means and standard deviations for these IAFs, 30 subjects were included in the 8.3hz condition, 31 subjects were included in the 12.5hz condition, and 32 subjects were included in the stationary condition.

### EEG response to flickering stimulation

To assess the effectiveness of our stimulation in synchronizing brain oscillations, we conducted FFT’s on the last 500ms of stimulation, again using Hamming windows and zero padding to a 0.1hz frequency resolution. Power and inter-trial coherence (ITC) were computed. Power was computed in the same way as was described in the previous section. Note that this measure of power reflects the power of all ongoing oscillatory activity, whether phase-locked to the stimulus or not. In order to asses phase-locked activity, we used ITC, which is a commonly-used measure of how consistently phase-locked oscillations at a given frequency are across trials (e.g. Makeig et al., 2002; Spaak et al., 2014; Keitel et al., 2019). This measure is independent of power and would be 1 if there was complete phase-locking across trials and 0 if there was none.

The same metrics were also computed for the first 500ms after the end of the last stimulation cycle to assess whether stimulus-evoked synchrony persisted after stimulation. Although there was not enough data to reliably examine post-stimulation synchrony in trials with the target stimulus omitted, we reasoned that the jittered SOA of the target would prevent any synchronous evoked activity from contaminating the analysis of all stimulation trials. Power and ITC were averaged across a bilateral group of 10 sensors was used for these analyses: (PO8 PO7, PO6, PO5, PO4, POz, O2, O1, Oz). This grouping of sensors was used in order to reduce noise, capture variability in the flicker-evoked response between subjects and is the area showing the most ITC in the average data. Note that for these analyses we focused on comparing the 8.3hz and 12.5hz conditions because a broadband increase in post-stimulation ITC was observed in the stationary condition, possibly due to a larger evoked response to the offset of the stimulus, see Figure 4.

### Testing for rhythmicity in performance

We also measured whether there was any rhythmic fluctuation in two-flash discrimination rates after stimulation (similar to Ronconi & Melcher, 2017). We first detrended each subject’s discrimination rate data (sorted by SOA). We then used a moving average of ±1 SOA bin to smooth each subject’s data and then averaged across subjects. We used the MATLAB curve-fitting toolbox to fit sine waves to this data using a nonlinear least squares fitting procedure. As mentioned in Ronconi & Melcher, 2017, using moving average smoothing in this way can lower the best fitting frequency compared to an FFT analysis. A starting point was set at 4hz for the search algorithm, and no further bounds were used. Note that although we only had 10 behavioral time-points, this was sufficient in Ronconi and Melcher (2017), experiment 2, to detect flicker-induced fluctuations in two-flash discrimination rates at 6.5hz, suggesting that we had sufficient resolution to detect fluctuations at our stimulation frequencies.

## Results

### Two-flash fusion thresholds vs. visual stimulation

There was a significant main effect of stimulation condition on fusion thresholds (*F*(3,78) = 5.939; *p* = 0.001). Post-hoc t-tests revealed that this main effect was driven by significantly lower thresholds in the stationary condition, compared to all other stimulation conditions (all *p*’s < 0.05). No other comparisons were significant. This suggests that two-flash fusion thresholds were not influenced by alpha-band flicker frequency, providing evidence that IAF was not entrained by visual flicker. Psychometric fits and thresholds are shown in Figure 2. A possible explanation for the improvement in thresholds in the stationary condition could be related to changes in perceived contrast between the background and the target induced by stationary stimulus after effects (see Discussion).

**Figure 2.**
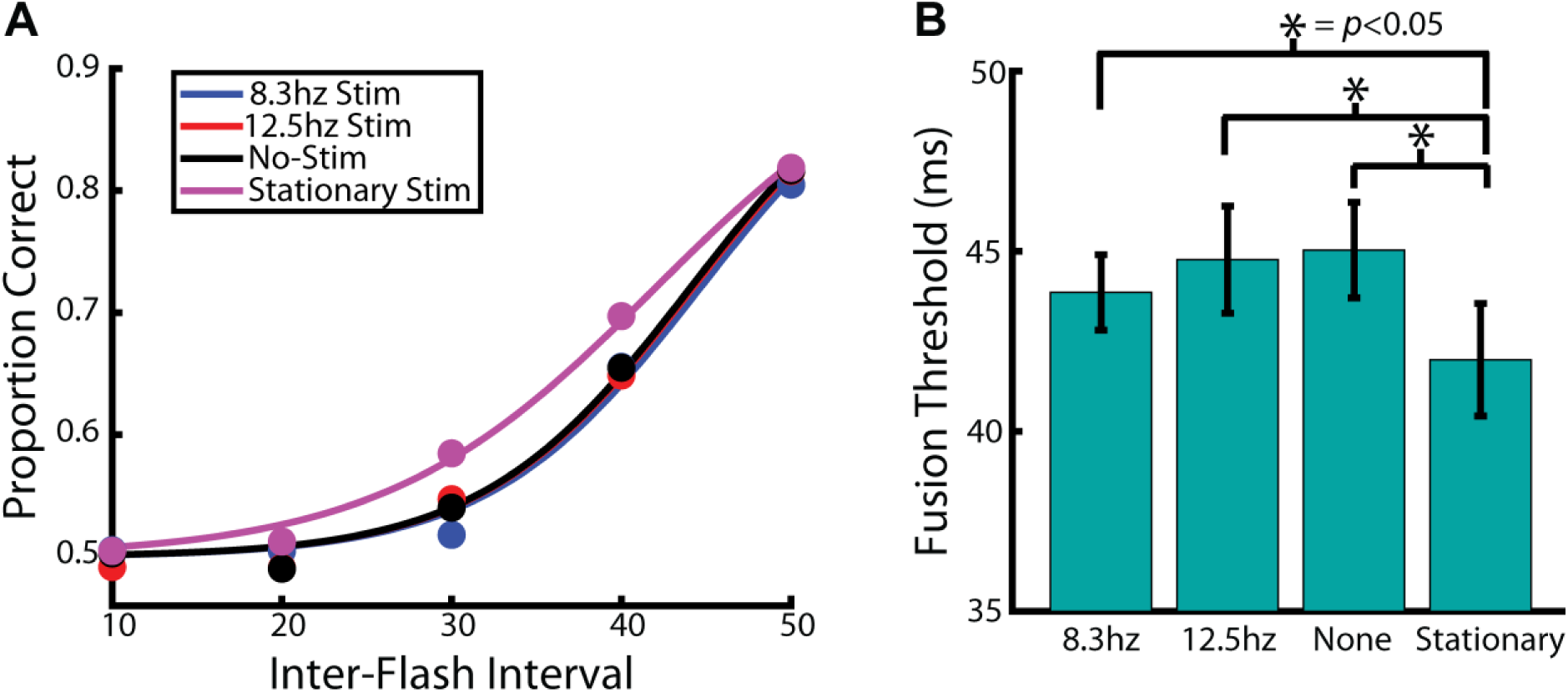
Task performance. A. Psychometric curves were fit to the two-flash discrimination performance, from which two-flash fusion thresholds were derived, shown in B. Fusion thresholds were significantly faster in the stationary condition compared to all other conditions. Error bars represent ± 1 SEM.

### Two-flash fusion thresholds vs. IAF

Replicating Samaha & Postle, 2015, two-flash fusion thresholds in the no-stimulation condition correlated with IAFs in both the eyes-closed and eyes-open resting sessions (eyes-closed: *rho* = −0.37; *p* = 0.037; eyes-open: *rho* = −0.43; *p* = 0.014). Since there was no effect of task condition on IAF during the ITI (*F*(2.18,74.14) = 1.806, *p* = 0.168, Greenhouse-Geisser corrected), one IAF was computed for the ITI across all trials. This set of IAF values was also correlated with two-flash fusion thresholds (*rho* = −0.36; *p* = 0.044). Eyes-closed IAF did not correlate with thresholds in any other task condition, see Figure 3. Eyes-open IAFs were marginally correlated with thresholds in the 12.5hz condition (*rho* = −0.34; *p* = 0.051) but not in the 8.3hz or stationary stimulation conditions. IAFs in the ITI did not correlate with thresholds in any of the stimulation conditions. See Table 1 for a list of all correlations.

**Table 1.** Two-flash fusion correlations for all IAF measurements and stimulation conditions.

|  | Eyes-Closed IAF | Eyes-Open IAF | ITI IAF |
| --- | --- | --- | --- |
| **8.3hz** | $\rho = -0.18$; $p = 0.308$ | $\rho = -0.24$; $p = 0.178$ | $\rho = -0.18$; $p = 0.311$ |
| **12.5hz** | $\rho = -0.26$; $p = 0.147$ | $\rho = -0.34$; $p = 0.051$ | $\rho = -0.25$; $p = 0.157$ |
| **Static** | $\rho = -0.08$; $p = 0.666$ | $\rho = -0.07$; $p = 0.712$ | $\rho = -0.12$; $p = 0.492$ |
| **No-Stimulation** | $\rho = -0.37$; $p = 0.037$ | $\rho = -0.43$; $p = 0.014$ | $\rho = -0.36$; $p = 0.044$ |

**Figure 3.**
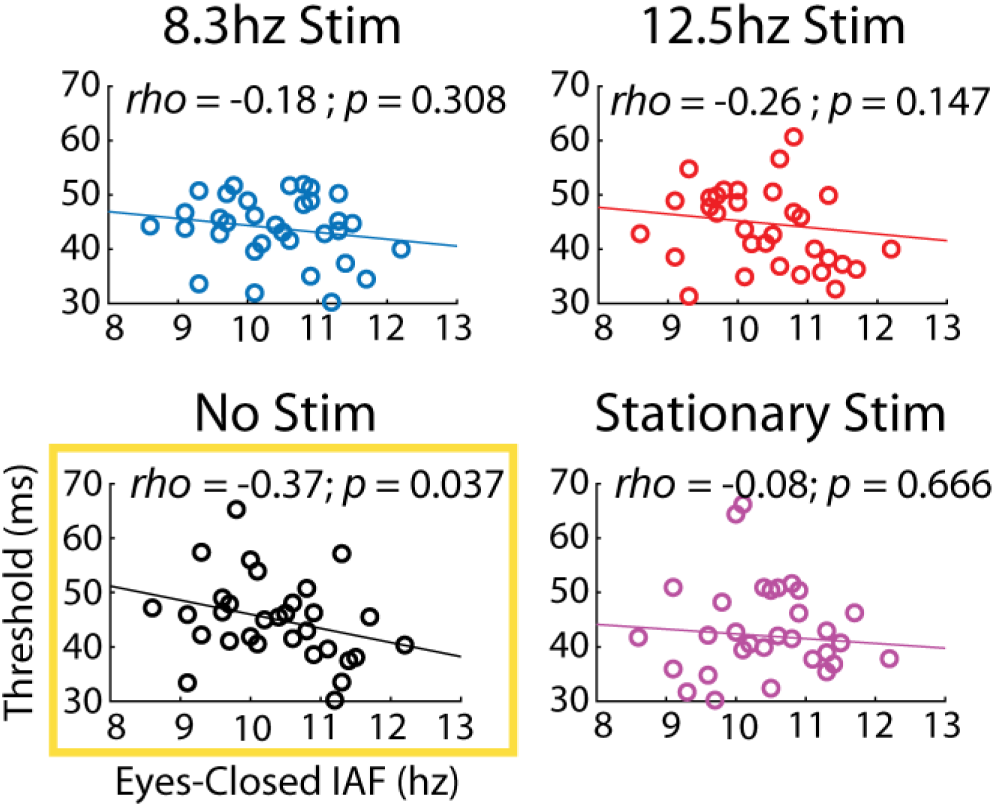
Correlations between eyes-closed IAF and fusion thresholds. The no stimulation condition was the only condition that correlated with IAF. See Table 1 for additional correlations between fusion thresholds and IAFs calculated from the eyes-open session and the ITI.

**Figure 4.**
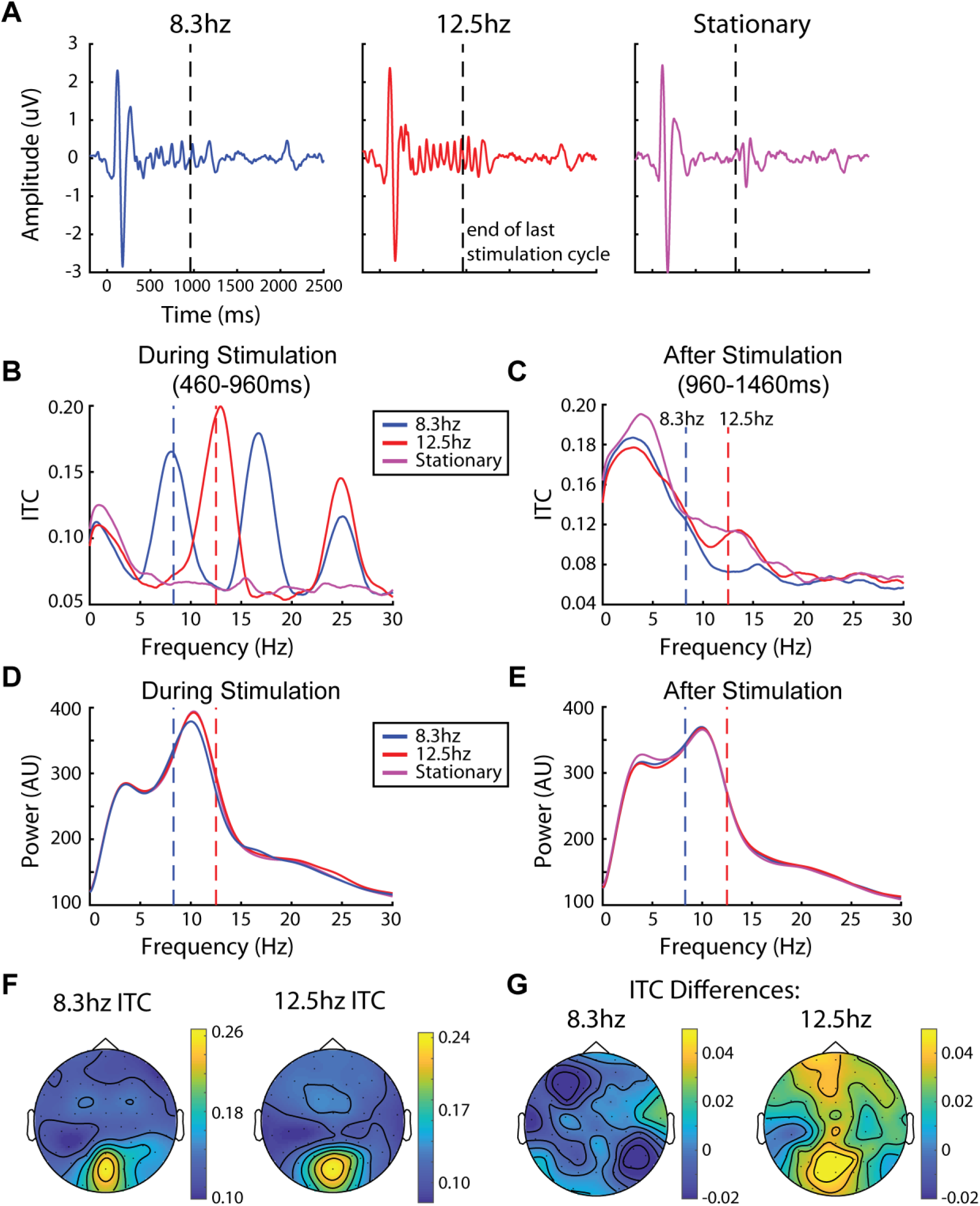
Task EEG data. All time and frequency plots are averaged from posterior sensors. A. Time-series of the stimulation conditions. Time 0 indicates the start of stimulation. Vertical dashed line indicates the end of the last flicker cycle. Note that this means that the entrainment stimulus turned dark-gray 40-60ms before this marker, see Methods. B. ITC during the final 500ms of stimulation. Vertical dashed lines indicate the stimulation frequencies. C. ITC in the 500ms after stimulation ended. D. Power during the final 500ms of stimulation. E. Power in the 500ms after stimulation ended. F. Topographies of the flicker evoked responses during stimulation. G. Difference topographies of the flicker evoked responses persisting after stimulation. The topography on the left is ITC at 8.3hz, subtracting the 12.5hz condition from the 8.3hz condition. The topography on the right is ITC at 12.5hz, subtracting the 8.3hz condition from the 12.5hz condition.

### Flicker-induced modulation of EEG

A strong oscillatory response was observed during both rhythmic stimulation conditions at their stimulation frequencies. This is visually apparent in the time domain as shown in Figure 4A. Statistically confirming this, ITC during the last 500ms of stimulation (shown in Figure 4B) was significantly enhanced in the rhythmic conditions at their stimulation frequencies compared to the stationary stimulation condition (8.3hz: *t*(35) = 7.31, *p* < 0.001; 12.5hz: *t*(35) = 7.56, *p* < 0.001). Despite strong phase-locking to the flickering stimuli, there was no effect of stimulation on IAFs measured during the last 500ms of stimulation (*F*(2,50) = 0.675, *p* = 0.514; mean IAF during 8.3hz stimulation = 10.43hz, SD = 0.89hz; mean IAF during 12.5hz stimulation: 10.45hz, SD = 0.87hz; mean IAF during stationary stimulation: 10.40hz, SD = 0.82hz).

As can be seen in Figure 4C, there were no clear peaks in the power domain at the stimulation frequencies, demonstrating that flicker-induced synchronization to our stimuli was primarily due to phase alignment. There was significantly greater power at 12.5hz in the 12.5hz condition compared to the 8.3hz condition (*t*(35) = 35.31, *p* < 0.001), but this was driven by an overall reduction in power in the 8.3hz condition over the full alpha range compared to the 12.5hz condition (8-13hz; *t*(35) = 22.1, *p* = 0.005). There was also no difference in power at 8.3hz (*t*(35) = 1.42, *p* = 0.101) compared to the 12.5hz condition.

Since it has been shown that individuals synchronize more strongly to stimuli that are flickered at rhythms closer to their IAF (Notbohm et al., 2016), we also measured whether there was a correlation between IAF and ITC during the last 500ms of stimulation. There were no correlations between resting or ITI IAFs and ITC in either flicker condition. These comparisons are shown in Table 2.

**Table 2.** IAF vs. ITC correlations.

|  | **Eyes-Closed IAF** | **Eyes-Open IAF** | **ITI IAF** |
| --- | --- | --- | --- |
| **8.3hz ITC** | $\rho = -0.02; p = 0.917$ | $\rho = 0.00; p = 0.989$ | $\rho = -0.05; p = 0.760$ |
| **12.5hz ITC** | $\rho = 0.11; p = 0.526$ | $\rho = -0.01; p = 0.971$ | $\rho = 0.11; p = 0.520$ |

Lastly, we tested whether there was flicker-evoked activity that persisted after the end of stimulation, as has been done previously (e.g. Spaak et al., 2014). ITC during the first 500ms after the final stimulation cycle is plotted in Figure 4D. Comparing ITC in the 8.3hz and 12.5hz conditions during this time window, we found that there was significantly greater ITC in the 12.5hz condition (*t*(35) = 3.97, *p* < 0.001) but not in the 8.3hz condition (*t*(35) = 0.85, *p* = 0.4). Post-stimulation ITC also appeared to be elevated from ∼1-5hz and ∼10-12hz in the stationary condition, which might be due to a slightly greater ERP to the offset of stimulation, which is somewhat visible in Figure 4A.

### Alpha frequency and power differences between resting sessions and the task

There was a significant change in IAF between resting sessions and the task (*F*(2,70) = 4.155, *p* = 0.020; eyes-closed mean IAF = 10.41hz, SD = 0.83hz; eyes-open mean IAF = 10.57hz, SD = 0.99hz; ITI mean IAF = 10.63hz; SD = 0.86hz). Follow up *t*-tests revealed that IAF was marginally higher during the eyes-open resting session compared to the eyes-closed session (*t*(35) = 1.97, *p* = 0.056), and IAFs in the ITI were significantly greater compared to eyes-closed IAFs (*t*(35) = 2.49, *p* = 0.018). This data is shown in Figure 5A-B.

**Figure 5.**
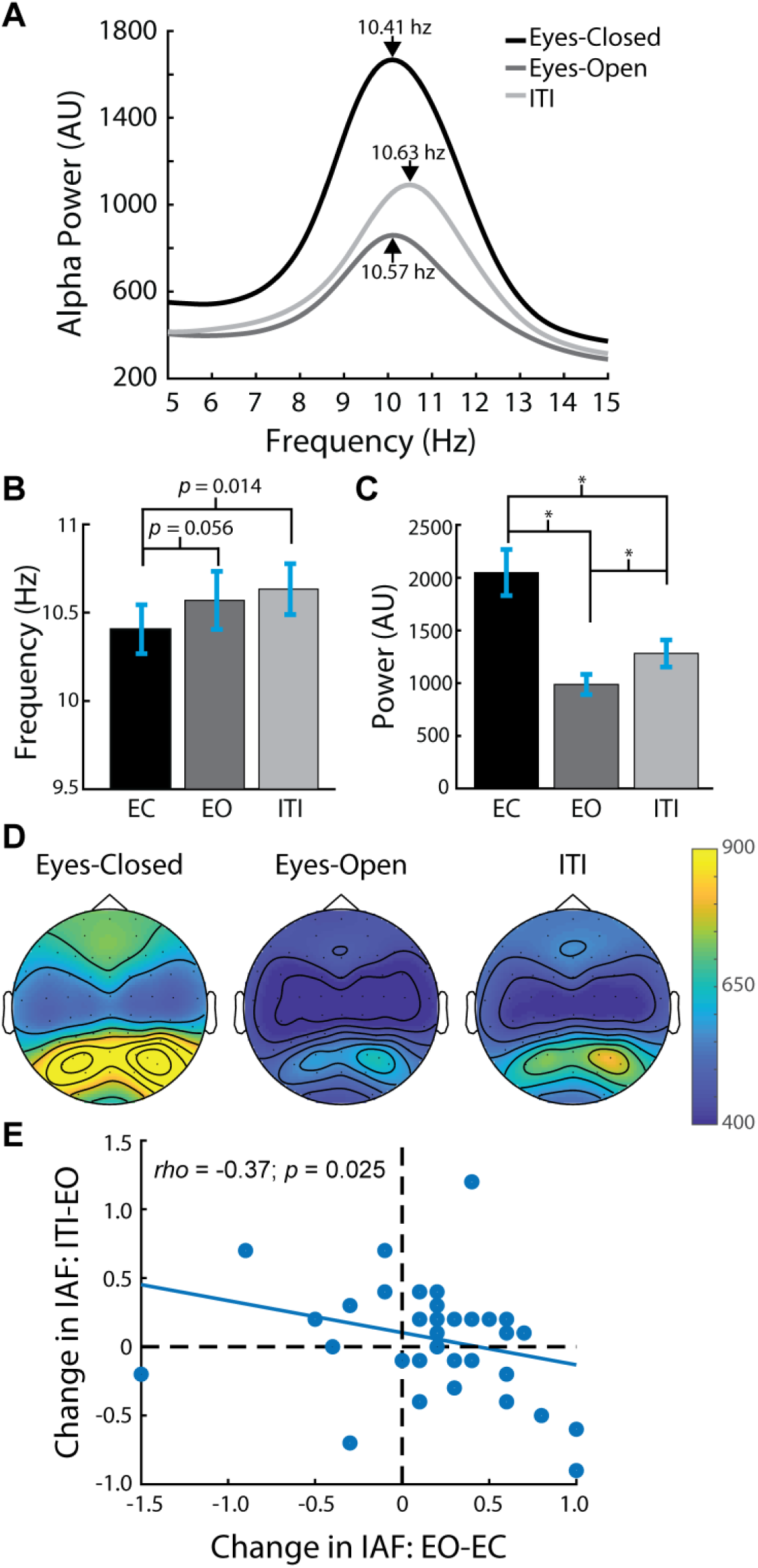
Resting and ITI alpha characteristics. A. Alpha power and peak frequencies at sensor PO6 (the sensor with maximum alpha across subjects) for the resting recordings and the ITI. B. IAFs for the resting recordings and the ITI (EC = eyes-closed; EO = eyes-open). C. Alpha power for the resting recordings and the ITI. * = p < 0.001. Error bars represent ± 1 SEM. D. Alpha power topographies for the resting recordings and the ITI. E. Scatterplot showing the correlation between the change in IAFs between EC to EO and EO to ITI.

Within subjects, IAFs were highly correlated between all recording sessions (eyes-closed vs. eyes-open: *rho* = 0.86; *p* < 0.001; eyes-closed vs. ITI: *rho* = 0.79; *p* < 0.001; eyes-open vs. ITI: *rho* = 0.91; *p* < 0.001). Interestingly, the change in IAF from the eyes-closed to the eyes-open session was negatively correlated with the change in IAF from the eyes-open session to the ITI (*rho* = −0.37; *p* = 0.025). This is shown in Figure 5E. These changes in IAF between sessions were not correlated with changes in alpha power between sessions (eyes-closed vs. eyes-open: *rho* = −0.20; *p* = 0.242; eyes-open vs. ITI: *rho* = −0.06; *p* = 0.748).

Alpha power also changed significantly between resting sessions and the task (*F*(2,70) = 27.954, *p* < 0.001). Follow up *t*-tests revealed that power was greater in the eyes-closed session compared to the eyes-open session (*t*(35) = 6.25, *p* < 0.001), and compared to the ITI (*t*(35) = 4.42, *p* < 0.001). Alpha power in the ITI was greater than in the eyes-open session (*t*(35) = 4.01, *p* < 0.001). See Figure 5C-D. Within subjects, alpha power was also highly correlated between all recording sessions (eyes-closed vs. eyes-open: *rho* = 0.70; *p* < 0.001; eyes-closed vs. ITI: *rho* = 0.91; *p* < 0.001; eyes-open vs. ITI: *rho* = 0.66; *p* < 0.001).

### Post-Experiment Questionnaire

Eyes-closed IAFs were not significantly correlated with any of the questions on the post-experiment questionnaire. Eyes-open and ITI IAFs were both significantly correlated with how much sleep participants reported usually getting: (eyes-open: *rho* = 0.36; *p* = 0.035; ITI: *rho* = 0.41; *p* = 0.015). No other correlations between the questionnaire items and IAF were significant.

### Rhythmic fluctuation of two-flash fusion discrimination

Although we observed stimulation-locked neural oscillations persisting after the flicker was over, we found that two-flash discrimination appeared to fluctuate at a general alpha frequency rather than at the rhythm of stimulation. Smoothed behavioral data and sinusoid fits are shown in Figure 6. Significant sinusoidal modulation of the aggregate behavioral data was found in the 12.5hz condition and the stationary conditions (stationary condition: *p* = 0.008, *adj-R*^*2*^ = 0.772, best-fitting frequency: 10.2hz; 12.5hz condition: *p* = 0.032, *adj-R*^*2*^ = 0.600, best-fitting frequency: 9.19hz). The 8.3hz condition did not exhibit significant rhythmic modulation (*p* = 0.225, *adj-R*^*2*^ = 0.313). We also fit the data in the no-stimulation condition by using arbitrary SOAs 100-190ms before two-flash presentation and as expected found no significant modulation (*p* = 0.865, *adj-R*^*2*^ = −0.332).

**Figure 6.**
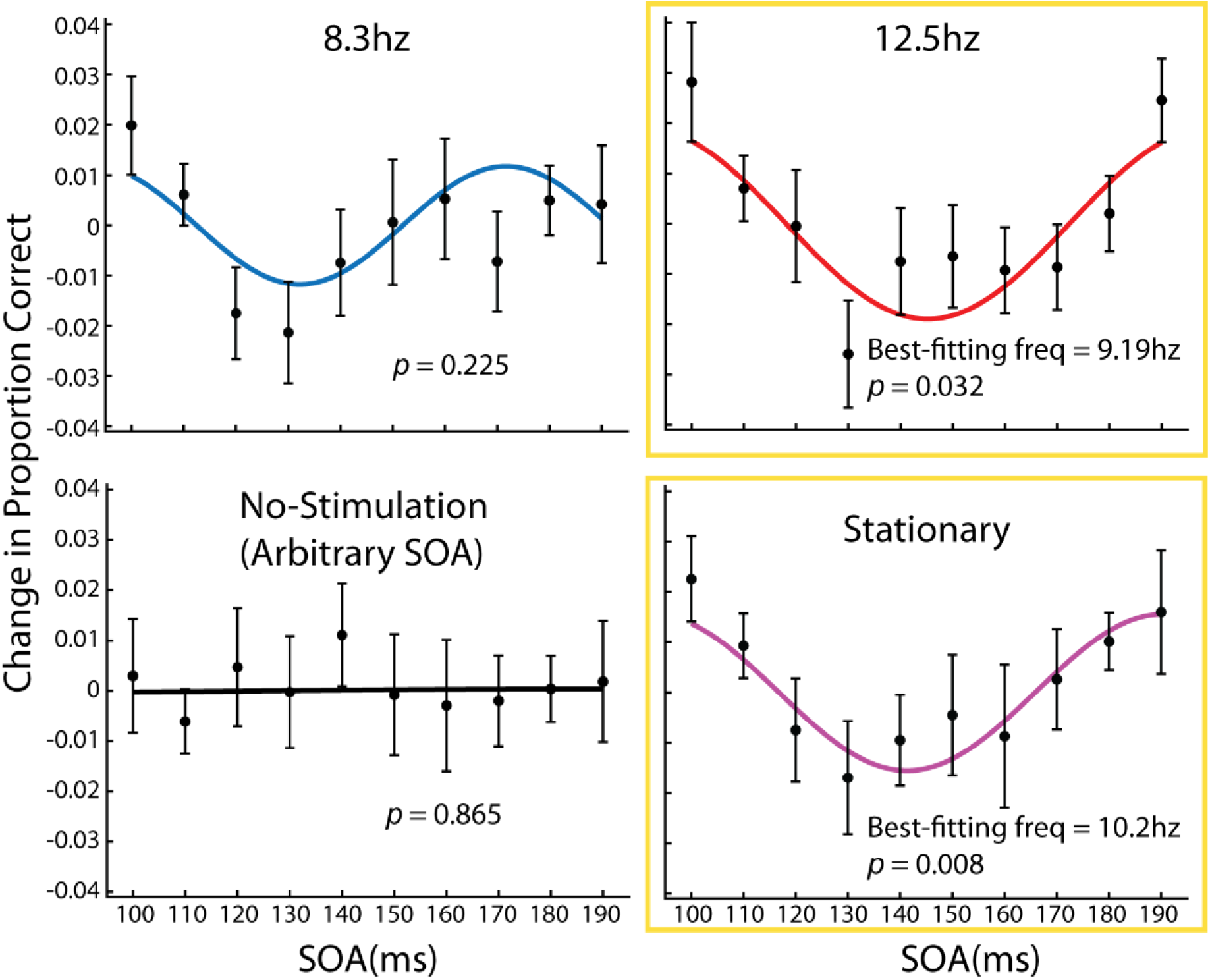
Rhythmic fluctuations in two-flash discrimination. Each plot contains the detrended and smoothed (via moving average) behavioral data points, error bars represent ±1 SEM. Also plotted are the best-fitting sinusoidal curves of the smoothed data. P-values were derived from random permutation testing, the fits were significant in the 12.5hz and stationary conditions.

## Discussion

### No evidence for the malleability of IAF

In this study, we found that rhythmic visual stimulation in the alpha-band did not cause changes in IAF, nor did it influence two-flash fusion thresholds. Importantly, there was a significant correlation between IAF and two-flash fusion thresholds in the no-stimulation condition, replicating Samaha and Postle (2015) and demonstrating that our task should have been sensitive to stimulus-induced changes in IAF persisting after stimulation. We also did not find a correlation between IAFs and the strength of synchronization at either stimulation frequency, which would also be expected if IAF were being entrained at sufficiently high intensity (Notbohm et al., 2016). These findings suggest that IAF is not malleable to bottom-up stimulation but instead reflects neural processes that regulate perception internally regardless of changes in the environment.

### Evidence for the entrainment of alpha activity independent of IAF

Previous studies have found that a short period of flicker at alpha frequencies can induce a corresponding fluctuation in perception at the flicker frequency (Mathewson et al., 2012; Spaak et al., 2014; Kizuk & Mathewson, 2016). Considering our null finding, this implies that the alpha activity which was entrained in these studies is separate from IAF. As was also found in these studies, we found stimulus-locked activity that persisted after the offset of the 12.5hz stimulus, supporting the notion that alpha entrainment involves activity that does not contribute to IAF.

Two other aspects of our data support this idea: 1) The topography of the post-stimulation synchrony was more posteriorly focused than that of the alpha oscillations measured during rest and the ITI. 2) Despite producing strong phase-locking, 8.3hz and 12.5hz flicker did not modulate ongoing power. This demonstrates a dissociation between measures of power and phase-locking which we were able to capitalize on to measure IAF and stimulus-locked entrainment separately. This analysis strategy has been recently emphasized by Keitel et al. (2019), who showed that attention to a lateralized stimulus flickering at 10hz or 12hz resulted in an increase in phase-locking and a decrease in power over contralateral scalp locations. These opposing effects similarly demonstrate the existence of two functionally-distinct populations of alpha activity over posterior areas.

### Phase-reset of alpha may have caused fluctuations in discrimination performance

Interestingly, two-flash discrimination performance after 12.5hz and stationary stimulation fluctuated at frequencies suggestive of a phase-reset of non-flicker-locked alpha activity at the end of stimulation. Perhaps this is the same alpha which contributes to IAF and two-flash fusion thresholds, which would again support the independence of IAF and entrained alpha oscillations. In support of the possibility of IAF phase-reset, a previous study found that after rhythmic stimulation at 10.6hz, within-subjects discrimination performance fluctuations correlated with their IAFs (de Graaf et al., 2013). On the group level, this manifested as a ∼10hz fluctuation in discrimination. After 5.3hz stimulation, a significant 10hz fluctuation in discrimination also occurred on the group level. Although within-subjects performance was not fitted for this condition, this could have also been the result of IAF being phase-reset by the final entrainment stimulus.

However, if the discrimination fluctuations in our data were due to IAF phase-reset, we would also expect the correlations between two-flash fusion thresholds and IAF to remain in at least the 12.5hz and stationary stimulation conditions. This was the case in one of the comparisons between IAFs and fusion thresholds in the 12.5hz condition (eyes-open IAFs were marginally correlated with fusion thresholds, p = 0.051), but in all other comparisons in all of the stimulation conditions, IAFs did not correlate with two-flash fusion thresholds. This could suggest that IAF was influenced in a non-systematic way or that there was a third variable influencing two-flash fusion discrimination thresholds after visual stimulation.

### IAF is influenced by cognitive factors

Although IAF was not systematically influenced by our stimuli, we did find that IAFs increased significantly during the ITI compared to the eyes-closed resting recordings and there was also a trend (p = 0.056) towards significantly greater IAF in the eyes-open condition compared to the eyes-closed condition. This is similar to another study which has shown an increase in IAF during a task compared to rest (Haegens et al., 2014). In their study they found that IAF was increased during an N-back working memory task compared to rest and passive visual stimulation. They found that IAF increased even more in the harder condition of the task, suggesting that this change was due to task engagement. The marginal increase we found in IAF from the eyes-closed to the eyes-closed session suggests that alertness may also influence peak alpha frequency.

That IAF can be influenced by both alertness and task engagement is consistent with an fMRI study which found correlations between power in the lower and upper portions of the alpha-band (7-10hz and 9-12hz, respectively) and two functional networks (Sadaghiani et al., 2010). More specifically, power in the lower alpha-band was negatively correlated with activity in the dorsal attention network, which was comprised of the intraparietal sulcus, the frontal eye fields, and the middle temporal cortex. Power in the upper alpha-band was positively correlated with a tonic alertness network, consisting of the dorsal anterior cingulate cortex, the insula, and the thalamus. Significant correlations with both functional networks were found in a similar right occipital-parietal region as the sensor we used for our IAF analyses.

It is thus conceivable that the IAF increase from eyes-closed to eyes-open could have been due to an increase in activity in this tonic alertness network, resulting in a shift in activity towards the upper alpha-band and an increase in IAF. The IAF increase from eyes-closed to the intertrial-interval may have additionally involved the dorsal attention network, resulting again in another shift of activity towards the upper alpha-band. During 12.5hz and stationary stimulation, there was more power in the alpha band. It’s possible that these conditions were more stimulating than the 8.3hz condition, resulting in greater activation of the tonic alertness network.

We also found a negative correlation between the change in IAF between the eyes-closed and eyes-open sessions and the change in IAF between the eyes-open session and the task. This might reflect differences in arousal regulation. For example, participants who exhibited relatively large increases in IAF from the eyes-closed to eyes-open session often had a subsequent decrease in IAF from eyes-open session to the task. These individuals may have been unable to sustain their alertness level during the task, resulting in a decrease in IAF. Further supporting the idea that IAF is related to arousal regulation, eyes-open and ITI IAFs were correlated with self-reported average sleep duration. Interestingly, this correlation was not found with self-reported sleep duration on the night prior to the experiment, suggesting that arousal regulation is more strongly determined by long-term sleep habits.

### Improved fusion thresholds after stationary stimulation

A puzzling aspect of our data was that fusion thresholds after stationary stimulation were significantly improved compared to all other conditions. We speculate that this effect may have been due to an after-effect of the stationary stimulus. It is possible that the light gray stationary stimulus produced a negative afterimage which made the subsequent dark gray annulus appear even darker. This in turn would increase the contrast of the two-flash stimulus and enhance its detectability. Supporting this possibility, we observed an even stronger improvement in fusion thresholds in an earlier pilot experiment in which our entrainer stimuli spatially overlapped with subsequent two-flash stimuli (N = 5; data not shown). Another possibility is that the offset of the stationary stimulus increased the predictability of the target’s appearance, while being overall less distracting than the flickering stimuli.

### Potential limitations

In Notbohm et al. (2016), very high stimulation intensities were needed to produce signatures of IAF entrainment. Thus, the lower stimulation intensity used in our experiment may have prevented us from observing evidence of a shift in IAF. It is worth noting, however, that previous studies finding fluctuations in perception after alpha-band flicker used relatively low stimulation intensities, comparable to our own stimuli (Mathewson et al., 2010; Mathewson et al., 2012; Spaak et al., 2014; Kizuk & Mathewson, 2016). This suggests that IAF entrainment as measured by Notbohm et al. (2016) may be different from the alpha entrainment observed by these other studies and our own.

Although we did find EEG evidence for entrainment after 12.5hz stimulation, we did not find such evidence after 8.3hz stimulation. We speculate that this may be an issue of signal strength. Previous studies have found that post-stimulation fluctuations in the EEG are very short-lived. In Spaak et al. (2014), stimulation-induced fluctuations at 10hz lasted ∼300ms after the last flicker. At 12.5hz stimulation (80ms cycles), almost 4 cycles occur within that time period. At 8.3hz stimulation (120ms cycles), less than 3 cycles occur within 300ms, reducing the power to detect elevated ITC in this condition. Additionally, we chose to be conservative in analyzing post-stimulation synchrony: we windowed our FFT analysis to the end of the final stimulation cycle, rather than to the presentation time of the final entrainer, as other entrainment studies have done (Mathewson et al., 2012; Spaak et al., 2014; Kizuk & Mathewson, 2016). This means that the entrainment stimulus was dark-gray for half a cycle (40 or 60ms) before our post-stimulation window began.

### Visual entrainment as a general temporal expectation mechanism

Although there has been a great deal of focus on the specific entrainment of alpha oscillations, studies have shown that oscillations across a wide range of frequencies can synchronize to visual rhythms and form temporal expectations (Rohenkohl et al., 2012; Cravo et al., 2013; Sokoliuk & VanRullen, 2016; Gray et al., 2015). In Cravo et al., 2013, discrimination of a Gabor stimulus within noise was improved when the target was embedded within a 2.5hz rhythmic stimulus stream compared to a matched irregular stimulus stream. The authors interpreted their finding as a temporal expectation effect driven by the entrainment of 2.5hz oscillations because the phase of oscillations at this frequency correlated with performance. Sokoliuk & VanRullen, 2016 found that the detection of a small, near-threshold target fluctuated at the rhythm of a concurrently presented, circle which was sinusoidally modulated at 5hz, IAF, and 15hz.

Ronconi and Melcher (2017) used a very similar paradigm to our own and found that two-flash detection rates fluctuated in-synch with audiovisual stimulation at 6.5hz, 8.5hz, 11.5hz, but not 14.5hz. This supports the notion that oscillations within and below the alpha-band can be entrained in the service of temporal expectation. We suspect that the same stimulation-induced fluctuations of two-flash detection rates were not found in our data because our task was not as sensitive to rhythmic temporal expectations. Target stimuli in their task spanned 56ms (6ms flashes with 44ms in-between) while target durations in our task were 90-130ms (40ms flashes with 10-50ms in-between), which made it less likely that performance would differ between phases of the entrainment rhythm. Ronconi and Melcher (2017) also found that two-flash discrimination was significantly better after 8.5hz stimulation compared to 11.5hz stimulation, which also differs from our data but is still opposite to what would be expected if IAF-specific neural populations were being entrained.

Considering that entrainment of oscillations and behavior occurs from delta to alpha frequencies in tasks involving rhythmic temporal expectations, it is likely to be a general mechanism not specific to oscillations in the alpha-band. Studies finding rhythmic fluctuations in behavior after alpha-band flicker might have been tapping in to this mechanism rather than entraining alpha-specific activity (e.g. IAF). Further work is needed to validate this idea by more thoroughly testing for flicker-induced fluctuations in perception coinciding with persistent flicker-locked oscillations at a wider range of stimulation frequencies.

